# Following the Robot’s Lead: Predicting Human and Robot Movement from EEG in a Motor Learning HRI Task

**DOI:** 10.1101/2025.01.08.629388

**Authors:** Tanaya Chatterjee, Adrien Guzzo, Alejandro Tlaie, Charalambos Papaxanthis, Jeremie Gaveau, Peter Ford Dominey

## Abstract

Human-robot interaction (HRI) offers unique opportunities to study the neuroscience of human motor control through controlled and reproducible sensory stimuli. In this study, we introduce an innovative neuroscience-HRI framework inspired by the Serial Reaction Time (SRT) task, that combines EEG with a task where a humanoid robot performs preprogrammed movement sequences that are mirrored by a human participant in real time. The use of a humanoid robot ensures precise and repeatable sensory-motor stimuli in the 3D peripersonal space of the participant, providing experimental conditions that may be challenging to replicate with traditional methods. Behavioral performance is assessed by measuring the temporal lag between human and robot movements, which decreases with training, reflecting motor sequence learning. Concurrently, EEG data from the human participant is analyzed to reveal neural correlates of learning and movement dynamics. Event-Related Spectral Perturbations (ERSP) in theta, mu, and beta frequency bands demonstrate distinct patterns associated with rest, fixation, and movement. Furthermore, the ERSP changes over successive trials reflect the progression of sequence learning, highlighting the relationship between neural oscillations and motor learning. A Markov-Switching Linear Regression model further decodes EEG signals to predict movement parameters including both human and robot position and velocity in a time-resolved manner. Our findings highlight the potential of HRI as a robust platform for neuroscience research and underscore the value of EEG-based neural decoding in understanding motor sequence learning. This work suggests further advances for integrating robotics into neuroscience and rehabilitation research.

## Introduction

Understanding the organization of sequential behavior is one of the most interesting and challenging question in human behavioral neuroscience (1, 2). Addressing this question,in 1987 Nissen and Bullemer (3) published a study of human sequence learning using a simple computer setup with visual stimuli and keyboard responses which served as the basis for a vast and productive field of research that continues today, known as the serial reaction time (SRT) task.

While the SRT task has been of major importance in understanding human motor sequence learning (4), it is often restricted to finger movements on a keyboard. Recent studies using SRT tasks with more extended planar arm movements (5, 6) and pointing movements (7) have demonstrated robust sequence learning. Such learning effects should also be revealed in terms of neurophysiological markers. In human motor control, synchronized neural activity has been consistently demonstrated to become desynchronized at movement onset. This event related desynchronization (ERD) reflects neuronal activity which is increased and asynchronous during movement preparation and execution, leading to a reduction in the amplitude of oscillatory activity within specific frequency bands (reviewed in (8)). During continuous hand movement, a sustained ERD was observed in Mu and Beta frequency ranges (9). Likewise, alpha and beta ERD/ERS are modulated with by movement onset, continuation and offset in repetitive hand movements (10). In an SRT task where participants acquired both implicit and explicit knowledge of the sequence, (11) observed a progressive increase in ERD at 10Hz which was maximal when the participants acquired explicit knowledge of the sequence. This corresponds to a decrease in ERSP (event related spectral perturbation) over the course of the successive blocks. SRT learning similarly produced a progressive reduction in theta band (4–7Hz) power over successive blocks, with an increase in power for a final random block, that was not observed in the alpha/mu (7–12Hz) or beta bands (12–20Hz) bands (12). Thus, SRT learning can be characterized by different profiles of ERSP in theta, mu and beta bands over the course of sequence learning. While these changes in ERSP can reflect learning behavior at the level of blocks sequential behavior, it is of interest to understand how the EEG signal is related to finer grained elements of sequential behavior. Hidden Markov models (HMMs) have been successfully used to decode sequence learning behavior from intracranial electrode recordings in human patients (13). We have recently used a Markov-Switching Linear Regression (MSLR) model to link ongoing behavior to internal cognitive states on a trial to trial basis (14). In the current research we use MSLR to link cognitive states defined by neural activity to diverse parameters of the human sequential movement. Importantly we bypass the need for repeatedly presenting identical trials and then doing extensive post-hoc averaging because the MSLR model yields a time resolved estimate of cognitive states that can be directly compared to continuous behavior. In fact, this is an imperative step in the development of innovative solutions that can restore or enhance human motor functions. This is particularly true for technologies that interface the brain with computers but is also relevant to situations where one needs to monitor how a patient responds to a treatment (15–18).

Here we explore the relation between continuous behavior and the EEG signal in a human motor sequence learning task which extends the SRT paradigm to full arm sequential pointing movements towards 4 spatial targets in a face-to-face human robot interaction task as illustrated in Fig. 1A. In this task, the robot performs continuous sequences of full arm pointing movements to 4 locations in the shared space between it and the human. The human participant is instructed to follow the motion of the robot’s hand with her own hand, mirroring the motion of the robot. In this setting, we measured the lag between the robot and human, which served as a proxy for reaction times. We observe classic learning-related reductions in lag over successive blocks, and increases in lag during blocks in which the spatial or temporal structure of the sequence was perturbed, which serves as an indicator of sequence learning. In this context we investigate the relations between brain activity as revealed in recorded EEG and multiple parameters of the sequence learning behavior. As illustrated in Figure 1A, participants are in a face-to-face interaction with the humanoid robot, following the sequence traced by the robot with their own hand. Both have reflective markers on their arms allowing high resolution motion capture, and the human is equipped with a portable EEG recording system. Panel B illustrates the aligned robot and human motion data. Task-related ERSP and EEG signals are illustrated in panels C and D.

**Fig. 1.**
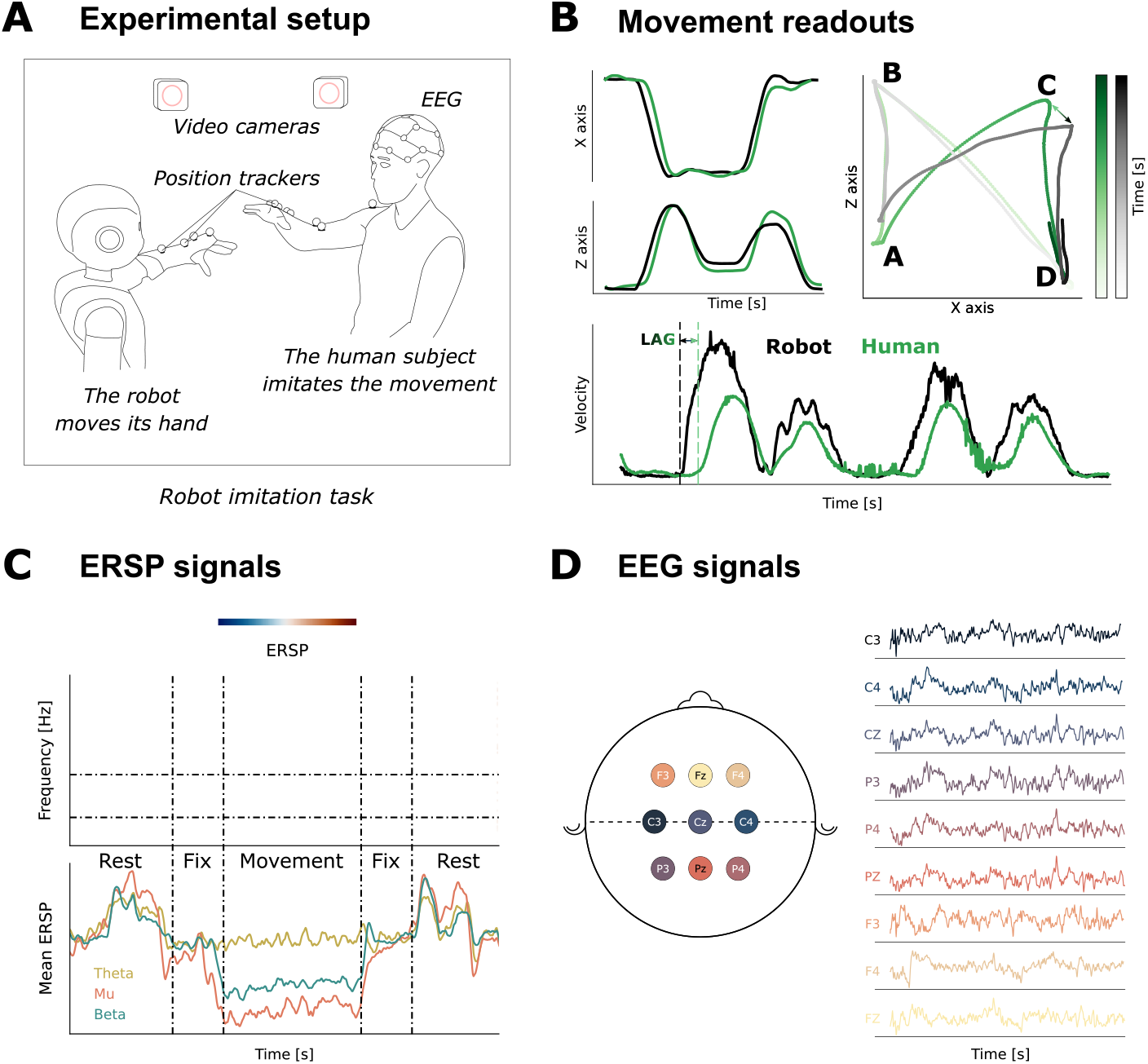
Recorded experimental data. A. Experimental setup : human participant face-to-face with Pepper humanoid. Robot performs preprogrammed motor sequences. Human mimics robot motion, mirroring the robot’s movements. Both have reflective markers for motion tracking. B. Motion tracking data : position in 1 and 2D, and velocity. Human in green, robot in black. Light-to-dark follows beginning to end in an example sequence of movements between the 4 spatial targets. C. ERSP grand average illustrating task-related desynchronization between task epochs of rest, fixation and movement for Theta, mu and alpha bands. Note strong desynchronization in mu and beta during the 80 element movement sequence. D. EEG signals mapped onto 10-20 coordinates for the 9 included electrodes that will be used in the MSLR model.

Using this neuroscience-HRI setup, this study makes several contributions to the field of motor sequence learning and human-robot interaction (HRI). First, by extending the classic Serial Reaction Time (SRT) task into a dynamic HRI context with a humanoid robot while recording simultaneous EEG, we create a novel platform that delivers precise, reproducible sensory stimuli, which is difficult to achieve with traditional non-robotic methods (6, 7). This advancement allows for a more naturalistic investigation of motor learning and underlying neural processes. Second, we provide new insights into the role of EEG in decoding motor behavior during real-time human-robot interaction, demonstrating that Event-Related Spectral Perturbations (ERSP) can be used to track the neural correlates of motor sequence learning across different task phases (8, 12). Lastly, by applying a Markov-Switching Linear Regression (MSLR) model, we show that neural activity can be decoded with high temporal resolution, allowing us to link EEG signals directly to behavioral parameters of the movement trajectory of both the human and robot. This approach builds on and enhances existing methods of neural decoding in motor control (16, 19), positioning HRI as a promising tool for further studies in neuroscience, joint action and adaptive robotics.

## Results

### Global ERSP modulation across Rest-Fixation-Movement

Figure 2 illustrates the ERSP modulation during the distinct task periods of Rest, fixation and movement. As shown in Figure 2A, we observed a clear effect for rest, fixation and the sequencing movement periods where Movement was most desynchronised and Rest periods were the most synchronised. In Figure 2B we see that these effects are modulated differently in the three frequency bands. At the first fixation (Fix1) and the experimental (Movement) periods, mu is more desynchronised than beta and theta. During the sequential movements (Exp), the three frequencies visibly diverge.

**Fig. 2.**
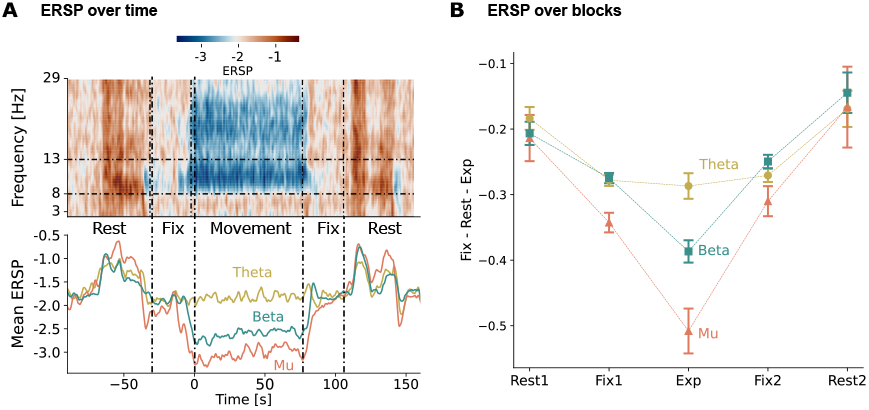
Main task-related effects on ERSP. **A**. ERSP values for the five main task-related periods averaged over all blocks and frequencies. ERSP is reduced for fixation vs rest, and for sequencing vs. Fixation. **B**. Frequency specific effects for the five task periods. ERSP reduction in Movement is greatest for mu as compared to theta andor beta.

These observations were confirmed by the 5 Period (Rest1, Fix1, Movement, Fix2, Rest2) *×* 3 Frequency ANOVA. We observed a main effect for Period (F (4, 68) = 26.57, p < 0.001, η^2^ = 0.325) and Frequency (F (2, 34) = 6.64, p = 0.004, η^2^ = 0.045). An interaction effect (F (8, 136) = 7.58, p < 0.001, η^2^ = 0.054) was also observed. The influence of the periods and the frequency were objectified through the post hoc comparisons with significant t-test and pholm values. During Fix1 Mu is significantly reduced with respect to Beta (p<0.05) and Theta (p<0.005). During Movement, Mu<Beta (p<0.05), Mu<Theta (p<0.001), and Beta>Theta (p<0.05).

### Sequence learning behavior

As illustrated in Figure 3, sequence learning can be characterized by a progressive reduction in the lag (which is a correlate of reaction time) between the human and the robot over the successive sequences. This is an index of learning that may also include sensorimotor learning independent of sequence-specific learning. To assess sequence-specific learning, we can observe the increase in lag in blocks 5 and 8 (perturbation blocks, see Methods), where the spatial or temporal structure of the sequence is modified, with respect to the preceding intact blocks 4 and 7. These learning effects were confirmed by a 2 × 2 repeated measures ANOVA. The dependant variables were the lags for blocks 4,5,7,8, where blocks 4 and 7 are training blocks, and 5 and 8 are perturbation blocks. The within subject variables were Epoch: (Early vs Late) and Type (Train vs Perturbation). There was a significant effect for epoch, F (1, 15) = 16.8, p < 0.001, as lags for blocks 4 and 5 were increased with respect to those for blocks 7 and 8, indicating overall lag reduction with successive blocks. We also observed a significant effect for type, F (1, 15) = 29.5, p < 0.001 which corresponds to the increased lags for the perturbation vs training blocks, which is the index for sequence specific learning.

**Fig. 3.**
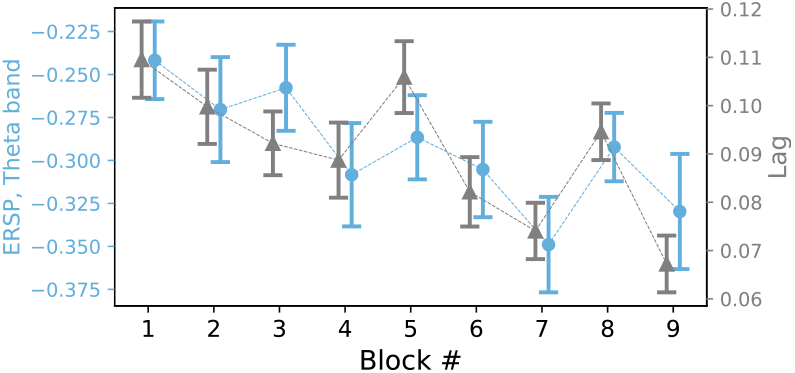
ERSP and Behavior. ERSP for all electrodes across 9 blocks compared with the behavioral measure of lag between the human and the robot. A significant reduction is observed in the ERSP (Theta) values of the first block (B1) versus the later training blocks B7, B9. A similar reduction is observed in the behavioral measures for lag.

### Relation between ERSP and Behavior

During the successive blocks of the sequence learning task, we observed that the temporal lag between the human and the robot was successively reduced. This lag reduction is a behavioral measure of sequence learning performance, and is illustrated in Figure 3, along with ERSP measures over the same successive blocks. There we can observe a correlation between the lag behavior and the ERSP values. Theta is correlated with lag (r > 0.8, p < 0.01), suggesting that theta oscillations may reflect apsects of the overall sensorimotor learning process.

### Learning-related effects on ERSP

Following the same logic as with behavioral results, ERSP values in different blocks can be compared to investigate the neurophysiological underpinnings of sensorimotor learning in the present task. Figure 3 illustrates the grand average ERSP values for theta and beta over the nine blocks, compared with the behavioural lag values. In terms of overall learning, we observe that the ERSP decreases progressively over the successive training blocks. These observations were confirmed by the 9 Blocks * 3 Frequencies ANOVA with main effects for Block (F (8, 136) = 5.82, p < 0.001, η^2^ = 0.062) and Frequency (F (2, 34) = 30.28, p < 0.001, η^2^ = 0.264), and the Block * Frequency interaction (F (16, 272) = 3.33, p < 0.001, η^2^ = 0.025). There was a clear difference between the first block (B1) and that of the later training blocks, B6(t = 4.286, pholm = 0.017), B7(t = 4.322, pholm = 0.017) and B9(t = 3.777, pholm = 0.050) where the later blocks had greater desynchronisation in ERSP. This reduction in ERSP over successive training blocks is an index of general learning. In order to verify that the specific sequence has been learned, we examine ERSP effects in response to perturbations of the sequence.

### Perturbation effects

We compared difference between two successive training blocks, vs. a training block followed by a perturbation block. We predicted that for two successive training blocks the ERSP will decrease (thus giving a negative difference), while for a training block followed by perturbation block, the ERSP will increase (thus giving a positive difference).

In the upper panel of Figure 4 we observe that in the theta band the perturbation effect is greater for temporal than for spatial perturbations. Temporal effet is significant (p<0.01) while Spatial is not(p=0.18). The lower panel of Figure 4 illustrates the electrode map with corresponding significance values for spatial, temporal and combined perturbations. These observations demonstrate that the ERSP is sensitive (1) to general learning over successive blocks, (2) to sequence specific learning revealed by responses to perturbations, and finally, (3) to the temporal vs spatial perturbations.

**Fig. 4.**
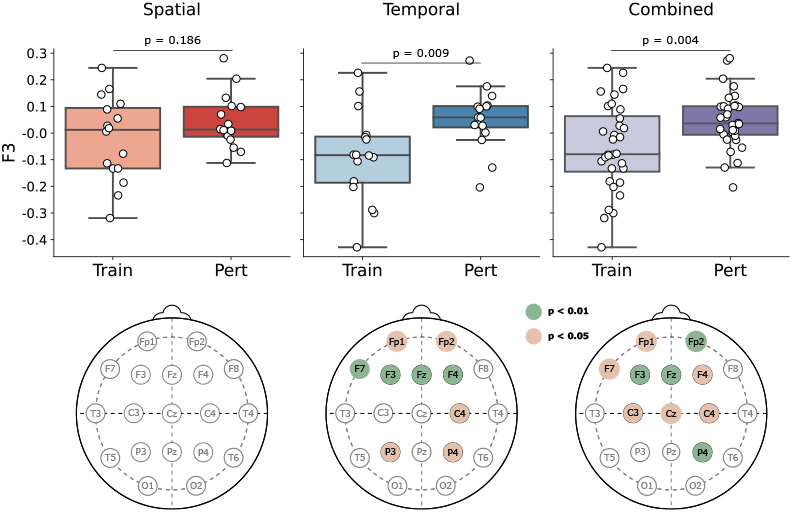
Effects of Spatial vs Temporal perturbations. Upper panel ERSP differences for Train-Train vs Perturbation-Train for Spatial and Temporal perturbations for electrode F3 in the theta band. Lower panels, scalp maps indicating electrodes with significant effects for spatial and temporal perturbation.

### Markov-Switching Linear Regression

Up until now, our analyses have focused on the block level. However, we hypothesized that the EEG signal contains information at a finer grained level. To test this hypothesis, we need a modelling approach that can recapitulate continous motion at an arbitrary timescale. We resort to the Markov-Switching Linear Regression (MSLR).

MSLR reflects the non-stationarity and regime shifts often present in behavioural data (14, 20–23), by flexibly accommodating complex temporal dynamics while keeping a relative simplicity, compared to deep learning-based methods (24). Moreover, the MSLR is less data-hungry than other common data-driven models (19, 25, 26). In our current case, the MSLR uses the EEG signal in each time point to predict a variety of behavioural readouts in the same time point, by assuming ‘hidden’ states. Each hidden state implies a different linear relation between values in the EEG signal and the corresponding movement features at the same time step. For instance, in one hidden state, the *lag* might be best predicted by the *F3* signal, while in another, the activity in *P4* might be most predictive. We used cross-validation to select the number of states (see below). For each trial, the model then outputs the predicted set of movement readouts as well as the probability of each hidden state. This functional flow is illustrated in (Fig. 5A). The model was trained and tested on data from all individuals.

**Fig. 5.**
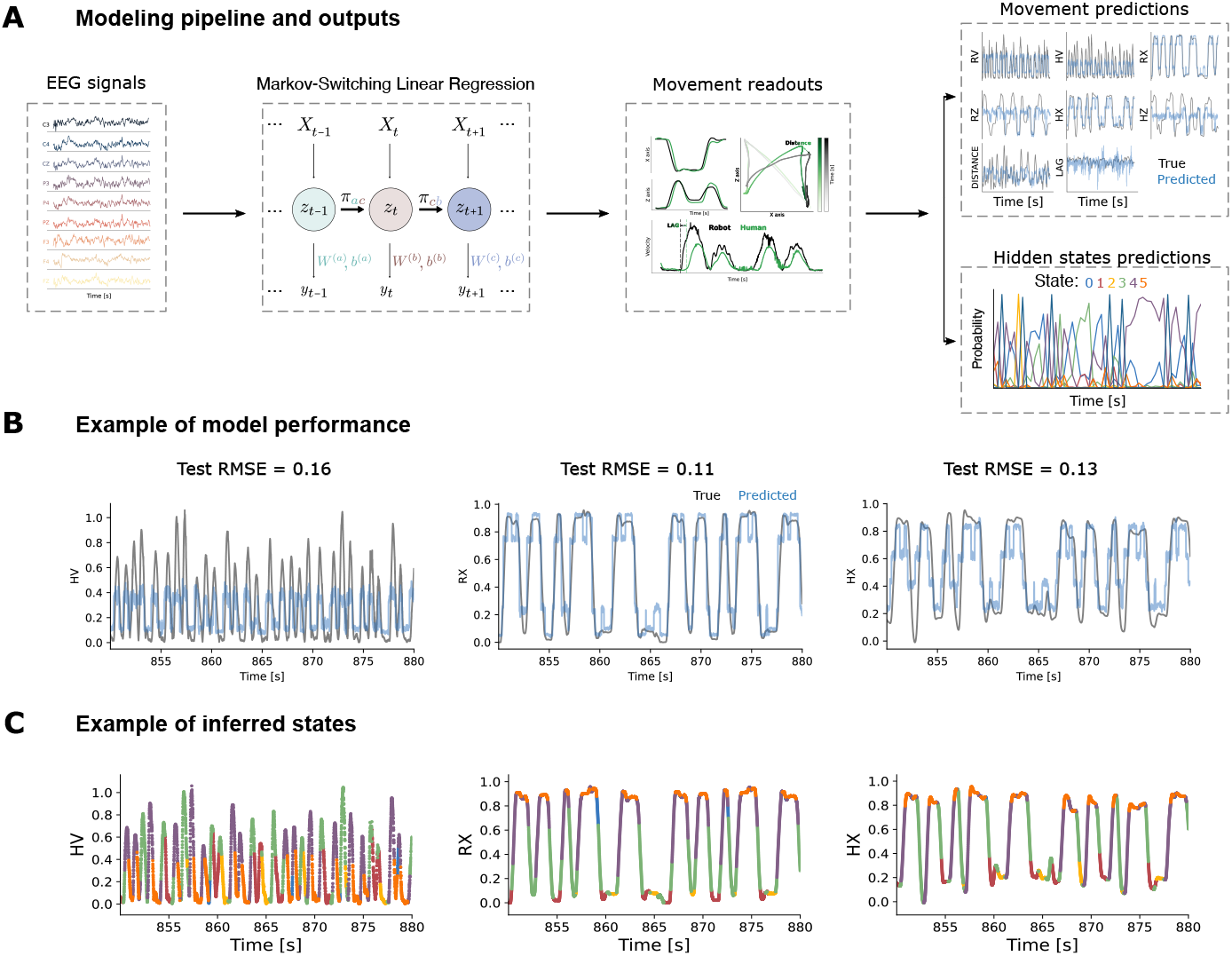
Modeling pipeline and example results for time-resolved predictions. **A)** The time-resolved model (Markov-Switching Linear Regression, MSLR) learns the linear mapping from EEG inputs to movement readouts. However, this linear relationship varies over time, through different hidden states. After training the model, it will output movement and hidden state predictions from novel EEG inputs. **B)** The model is able to predict human velocity (HV), robot X (RX) and human X (HX) positions; ground truth traces are shown in grey, model predictions in light blue. **C)** Mapping inferred states as color codes onto the predicted movement readouts, over time.

Mathematically, this model takes the form:

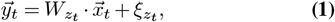

where 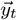 is the multivariate behavioural readout at time t,consisting on: vertical and horizontal movements (for the robot and for the human), their corresponding velocities, the lag of the human response and the distance between the robot and the human; z_*t*_ is the inferred state at time 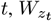 are the regression weights for state 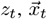 is the vector of EEG signals at time t, and 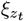 is a zero-mean Gaussian noise with variance 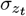.

To test if this approach was appropriate for our behavioural recordings, we quantified model performance for a different number of hidden states (n_*s*_) - which is the main free parameter of the MSLR. Model performance was tested by using time-blocked cross-validation (see Methods - *Model tuning*). We report (Fig. S1) that the cross-validated RMSE decreases as n_*s*_ increases. Since the error rate of model predictions began to saturate with increased model complexity, we decided to use n_*s*_ = 6 as a compromise between model complexity and performance.

### Decoding Movement from the EEG signal

Once the optimal number of hidden states was established, we performed tests on held-out data which showed a similar performance (Fig. 5, B) and (Fig. S2, B), indicating that the high predictive performance was not due to overfitting. There we see the clear superposition of the movement predicted from the EEG and the actual movements, with corresponding RMSE values. We highlight that these results are non-trivial, as the model was trained on finely sampled data; this is even more important when inspecting the actual structure of the time series: there are sudden variations, so the model had to predict these with a high level of precision (see Fig. 5 B for a zoomed-in example) or, otherwise, the RMSE would dramatically increase; in other words, for the RMSE to be small in such a finely sampled time-series, not only does the amplitude matter but also the time alignment. Jointly, this first set of results indicate that EEG recordings can indeed simultaneously predict a variety of ongoing movement readouts at the level of individual movements. These results also suggest that EEG and movement have a fluctuating relation over time, as reflected by the multiple underlying states.

To explore if the hidden states showed attributes that could be reflective of internal planning states, we first characterized their temporal dynamics. To this end, we examined the temporal structure of state transitions. The state transition matrices, which show how likely a timestep of a given hidden state is followed by a timestep of any (other or same) state (Methods - *Markov-Switching Linear Regression*), unsurprisingly revealed high values along the diagonal (Fig. S4A). This is because, as states are stable in time, self-transitions are much more common than cross-state transitions. When removing the diagonal and re-normalizing (row-wise), we find a set of more interesting results. We report (Fig. S4B) that there are bridging states (such as 3 → 0, 1, 2, 4), bi-directionally preferred (0 ↔ 5) and unlikely (2 ↔ 0) state transitions. This transition structure might in fact reflect the time structure of the task, as we will discuss next.

### Hidden states as performance states

As aforementioned, the MSLR outputs state probabilities per time step (Fig. S3A). Thus, we reasoned that if these states are reflective of ongoing movement (Fig. S3B), we could use their probabilities to classify actual movement categories. Note, however, that the model is agnostic to these categories: it was trained to predict the fine movement time traces, but it has no access to the way in which these movements are categorized. As we show in Fig. S3C, the inferred hidden states were consistently correlated with specific movement types (note that time points are colored in a time-cohesive manner).

Our rational was the following: if these inferred states are meaningful of movement throughout the task, we could use their inferred probabilities (Fig. 6 A, right) to decode the movement type (Fig. 6A, left). To do that, we trained a simple linear decoder and, per timestep, classify which movement type took place (out of the 19 possible categories, see the heatmap columns in Fig. 6C). We show (Fig. 6B) that we can predict these movement transitions significantly better than chance level (lower cross-entropy is better; Mann-Whitney U-test, p = 0.03); in this case, the surrogate distribution is built by shuffling the ground-truth labels and computing the cross-entropy 100 times. Note that the model was never exposed to task position labels (“A”, “B”, etc), but just to the raw time traces of movement readouts. Hence, this result highlights that the model is indeed picking up on time information and that hidden states can be traced back to the task structure, as we shall show in the next section. Furthermore, when we inspect the state importance (as captured by the decoder weights, Fig. 6C), we report that transitions involving *Rest* and location *D* have the most significant effects, with *Rest-D* greatly increasing state importance in State 3 and 4 and drastically decreasing in State 5. For instance, whenever the model inferred a high probability of state 2, the participant was most likely to move initiating at the right-hand side (*C-D, D, D-A, D-B, D-C* or *rest-D*), while those periods with a high probability of being in state 5 were most likely to move initiating at the left-hand side (*A-B, A-C, B, B-A, B-C, C, C-A* or *C-B*). These findings suggest that the influence of spatial movements on state dynamics is highly state-dependent, with specific states playing a pivotal role in predicting different fine transitions. The analysis underscores the complex interplay between spatial transitions and state importance, providing insights into how state dynamics can differentially disambiguate between fine movement types.

**Fig. 6.**
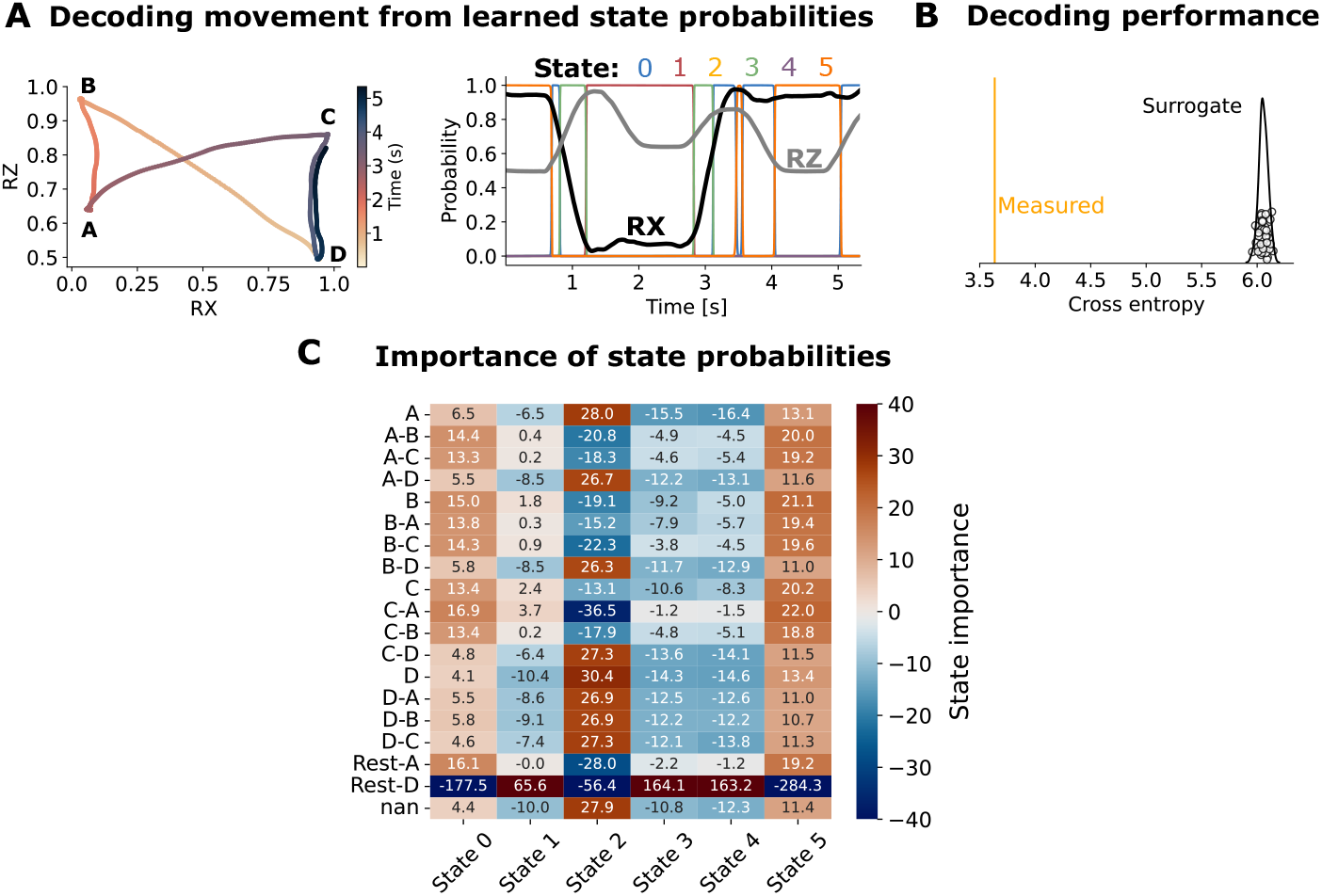
Using the learned state probabilities to decode movement type. **A)** Example 2*D* movement (left) for the robot (*x* and *z* coordinates). Dots are colored with respect to their occurrence in time. We also show that same time window unrolled over time (right), where state probabilities are the predictors to decode movement type. Movement transitions (RX or RZ) have its transition counterpart in state probabilities. These transitions in state probabilities are sharp both due to the model high certainty and to the high sampling rate of the input data. **B)** The linear decoder is significantly below chance level for the cross-entropy (lower is better). Given that the model was never exposed to task position labels (“A”, “B”, etc), but just to the raw time traces of movement readouts, this result highlights that the model is indeed picking up on time information and that hidden states can be traced back to the task structure. The surrogate distribution is built by shuffling the correct labels and computing the cross-entropy 1000 times. **C)** State importance, as measured by the weights of the linear decoder. There seem to be 3 main types: states that are useful to predict left-initiating movement (state 1), states that are useful to predict mostly right-initiating movements (states 0 and 5) states that were not useful to predict anything but *Rest-D* (states 3 and 4).

### Interpreting hidden states

Now that we have shown that these hidden states are useful to predict fine-grained movement and to classify movement type, we set out to shed some light on what these states correspond to. In order to do that, we followed a reverse-engineering approach, as shown in Fig. 7. We show the average value of each movement readout, split by state; this leads us to subsequently label these states as:

**Fig. 7.**
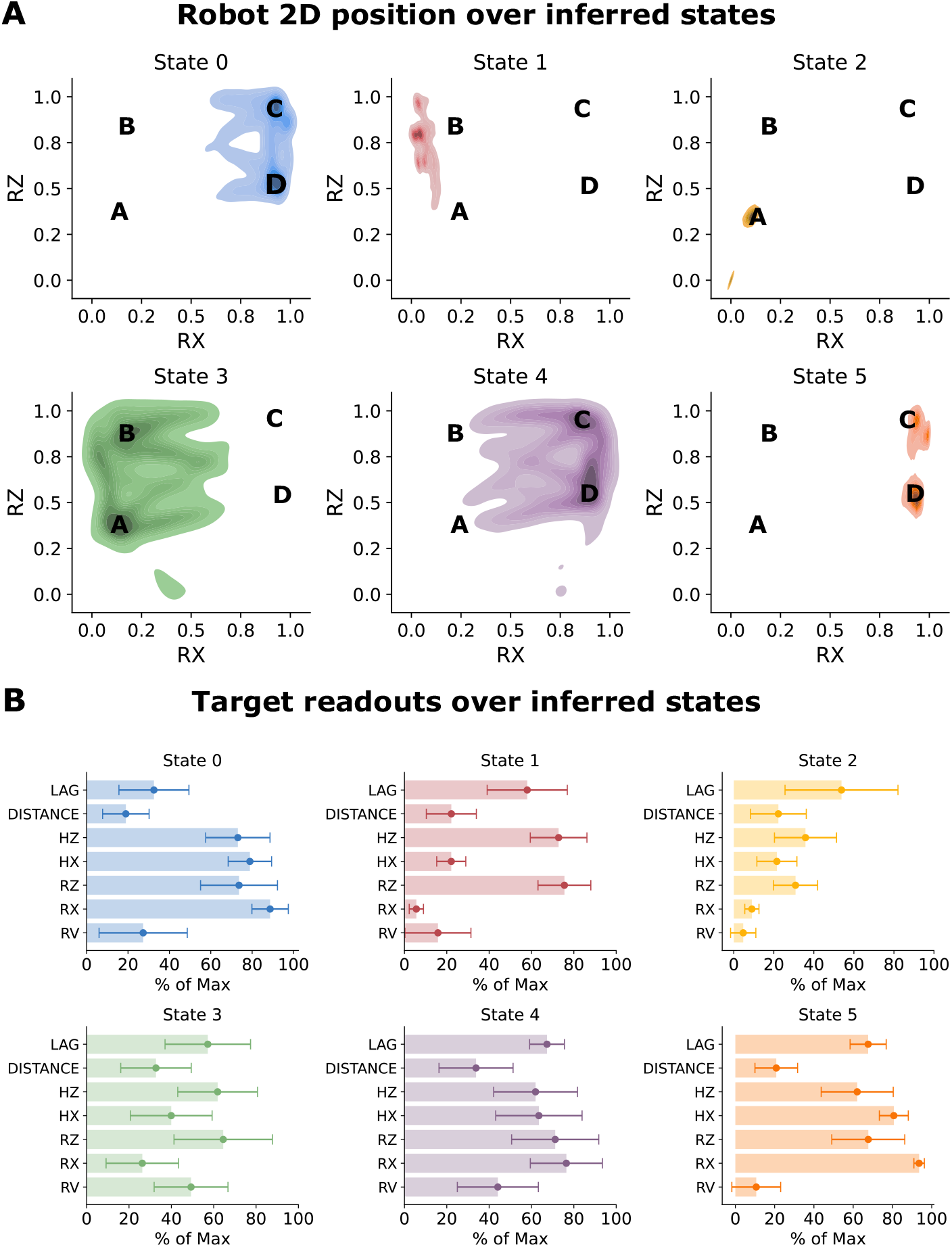
Looking at readouts to interpret hidden states. A) There is a clear state specialization, based either on movement (States 0, 3 and 4) or position (States 1, 2 and 5). Averaging target readouts, conditioning on inferred states, allows us to enhance state interpretability.

- State 0: right-hand side, engaged (low spatial variance, right-hand side movement, low lag, low distance, mid R_*v*_).
- State 1: B-location, movement (low spatial variance, left-hand side movement, high lag, low distance, high lag, mid distance, mid R_*v*_).
- State 2: A-location, movement (low spatial variance, mid lag, small distance, low R_*v*_.
- State 3: left-hand side, transition (high spatial variance, left-hand side movement, low precision, high distance, high lag, high R_*v*_).
- State 4: right-hand side, transition (high spatial variance, right-hand side movement, low precision, highest distance, highest lag, high R_*v*_).
- State 5: right-hand side, engaged (low spatial variance, right-hand side movement, high precision, high lag, small distance, small R_*v*_).

These characterisations allow us to reconstruct a potential way in which states take place over time, in general. One likely sequence from the transition matrix in Fig. S4B is: 0 → 3 → 1 → 3 → 4 → 5 … This state sequence corresponds to the overall task structure: from right (spatially localised, low lag) to left (spatially non-localised), to left (spatially localised), to left (location transition, non-localised), to right (non-localised), to right (localised).

### Electrode importance

Now that we have shown that the EEG signal can accurately predict ongoing movement readouts and that we have successfully characterised the inferred states, we turn our attention to inspecting how each electrode contributes to model performance. It is important to note that we can take this step because of the high interpretability of the MSLR, as disentangling the way in which each predictor aids model prediction would be much harder with other types of models such as Gaussian Processes (27) or almost any deep learning-based method (28).

As the MSLR is a set of linear regressions, we can just inspect the weights of each predictor as a measure of its importance for predicting each feature. In this case, of course, each predictor has a set of weights, of shape (n_*mov*_, n_*s*_), where n_*mov*_ is the number of movement readouts we want to predict and n_*s*_ is the number of hidden states (Fig. S5 shows these matrices per electrode). Therein, we see that each electrode contributes differently to model performance, over hidden states. If we now collapse one of the dimensions (averaging over states), and rearrange over readouts (Fig. 8A), we find that there are similar patterns of electrode contributions (Fig. 8B), as measured by the cosine similarity between average weight vectors. We also report that human-human variables are more extremely (anti-)correlated than robot-robot ones. Particularly, if we inspect some of the most extreme overall (anti-)correlations (yellow boxes in Fig. 8B), we see (Fig. 8C) that the spatial distribution of these electrode weights is indeed very similar.

**Fig. 8.**
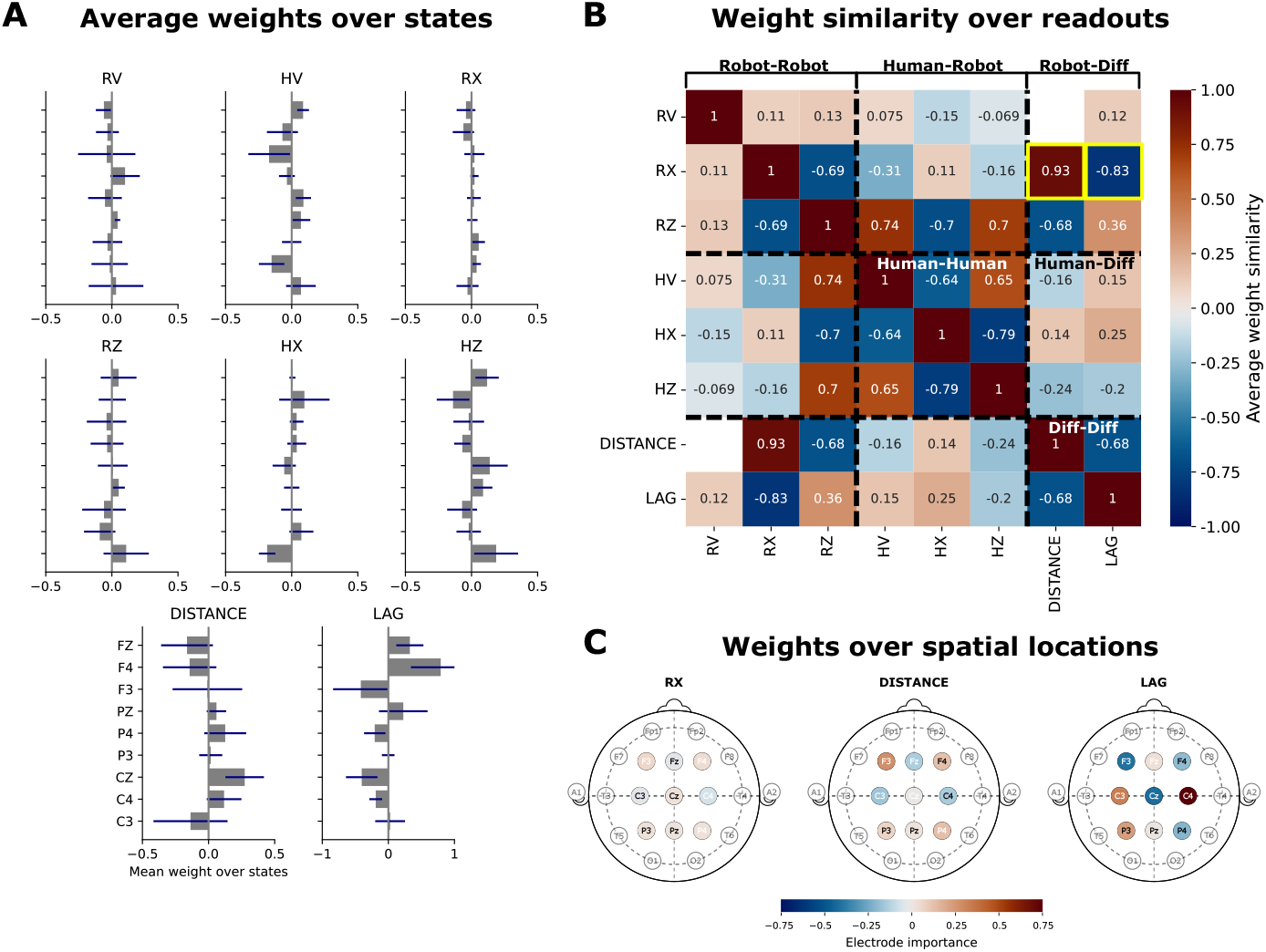
Electrode importance. A) Average learned electrode weights (gray bars) and standard deviation (blue lines) over states, for different behavioural readouts. We report different profiles for these average weights, depending on the to-be-predicted readout. B) Weight similarity over readouts. C) For three selected readouts (shown as yellow squares in B)), we show the actual scalp distribution of these weights, to show what the aforementioned similarity looks like.

## Discussion

This study introduces a novel human-robot interaction (HRI) paradigm that combines precise robotic stimuli with EEG-based neural decoding to investigate motor sequence learning.

### Advancing the State of the Art

Prior research on motor sequence learning has often relied on traditional paradigms, such as the Serial Reaction Time (SRT) task using computer-based visual stimuli and keyboard responses (3, 4). While these approaches have elucidated fundamental principles of motor learning, they often lack the ecological validity and dynamic interactivity found in real-world tasks. Recent extensions of the SRT task to more complex movements (e.g., arm-reaching tasks) (5–7) have demonstrated SRT learning with more realistic movment. At the same time, EEG has been used to characterize classic SRT learning (11, 12) in terms of learning-related modulation of oscillatory activity.

Building on this foundational work, our study employs a humanoid robot to deliver controlled and reproducible motor stimuli in the 3D peripersonal space of the participant. This approach addresses the challenge of creating consistent sensory inputs while enabling naturalistic interaction. Similar to recent advances in EEG decoding for motor control (16, 18), we leverage EEG to investigate the relationship between neural oscillations and continuous movement parameters. However, the current study goes beyond previous work by demonstrating that changes in Event-Related Spectral Perturbations (ERSP) reflect not only motor execution but also sequence-specific learning. Specifically, our findings reveal progressive reductions in theta-band power during training, consistent with the development of sensorimotor predictions, as well as increased theta activity during perturbation blocks, marking a neural response to disrupted expectations.

### Hidden Statesand Transition Dynamics

An important finding of this study is the identification of distinct hidden states in the EEG data, as modeled using Markov-Switching Linear Regression (MSLR). These states reflect different phases of the motor sequence learning process, and allow for decoding of the robot and human moveùents as illustrated in Figure 5. As shown in Figure 6, we identified three main types of states: (1) states that are predictive of left-initiating movements (state 1), (2) states that are predictive of right-initiating movements (states 0 and 5), and (3) states that were less useful for predicting specific movements but instead primarily associated with rest periods (states 3 and 4). Interestingly, states 3 and 4, which were less directly associated with specific movements, appear to play an important role in transitions between left and rightside movements. In Figure 5, we observe that these states are frequently visited during movement transitions, but only for short durations, highlighting their function as “transition states.” This is further confirmed in Supplementary Figure 3, where the temporal structure of these states aligns with short, transient periods of activity during the execution of movement transitions. These findings suggest that states 3 and 4, though not strongly tied to the 19 specific movement categories, are crucial for capturing the dynamics of movement transitions between different phases of the task.

### Contributions to Neuroscience and HRI

From a neuroscience perspective, this study contributes to a growing body of literature on the neural dynamics of motor learning. Our findings align with evidence that mu and beta desynchronization reflect movement execution (8, 9). However, by linking these oscillatory dynamics to behavioral metrics such as lag and trajectory, this research provides a more nuanced understanding of how neural oscillations support adaptive motor behavior. For HRI, this work highlights the potential of EEG-based neural decoding to enhance interactive systems. Similar to research on brain-machine interfaces for robotic control (15, 16), 2019), our findings demonstrate that neural signals can predict movement parameters in real time. The use of a humanoid robot adds an additional layer of innovation, as it creates ecologically valid interaction scenarios that are difficult to replicate with non-embodied systems. These insights could inform the design of adaptive robotic systems for rehabilitation, training, and beyond.

### Decoding human and robot movement in a joint action task

Interestingly, our task can be considered a form of coordinated joint action, in which the participant must closely observe the robot partner in order to mimic its movements (29). Extensive research has demonstrated that in the context of joint action, the participant’s behavior and brain activity is modulated by their own action, and their observation of the partner’s action (30). Indeed, related research (31) demonstrated that the P3a and P3b and the motor part of the contingent negative variation in the EEG signal were modulated by joint action, that is by the particpant’s actions and the partnter’s actions. We see the correlate of this in our decoding results. The MSLR states can be used to decode the spatiotemporal trajectories of motion, both for the human and the robot. This indicates that during this task, the participant’s EEG reflects not only their own movements, but those of the robotic partner as well. Such explicit distinct coding of one’s own actions and those of the partner should be of value in coordinating joint action (32).

### Limitations and Future Directions

While this study provides novel insights, there are limitations that warrant further exploration. This research utilized a single humanoid robot; future studies could investigate whether the findings generalize to different robot morphologies or interaction modalities. Moreover, while the MSLR model successfully decoded dynamic state transitions, its use could be complemented by other methods, such as deep learning or hybrid approaches, to explore additional latent structures in the neural data (19)(Keshtkaran et al., 2022). Expanding this framework to include multimodal data, such as eye tracking or electromyography (EMG), could provide a more holistic view of sensorimotor integration. Furthermore, incorporating bidirectional HRI—where the robot adapts its movements based on real-time neural feedback—could open new possibilities for studying and enhancing motor learning.

## Conclusion

By integrating neural decoding with behavioral analysis, this study builds on prior work and provides a comprehensive view of motor sequence learning in human-robot interaction. The reproducibility and precision of the robotic stimuli, combined with the dynamic insights from EEG data, underscore the potential of HRI as a tool for neuroscience and rehabilitation research. These findings lay the groundwork for future explorations into adaptive robotics, rehabilitation, and braincomputer interface technologies.

## Methods

### Participants

18 right-handed healthy subjects participated in the study (Mean age = 29.5 *±* 4.95, nine male). The participants were recruited through University’s internal email system. The demography consisted of master’s and PhD students and post doc researchers associated to the University of Bourgogne Franche-Comte (UBFC), France. The participants reported no history of neurological or psychiatric disorders. The study was carried out following legal requirements and international norms (Declaration of Helsinki, 1964) and approved by the ethics committee of the Université Bourgogne Franche-Comté (CERUBFC-2023-11-13_050).

### Apparatus

For the stimulus presentation of the motor task, we used an interactive humanoid robot Pepper controlled by the Choregraphe software (Aldebaran/United Robotics). Behavioral data were collected using a high fidelity motion capture system (Vicon) with markers placed on the human and robot arms, as illustrated in the methods Figure 4A. In the current paper we focus on the electrophysiological findings of the experiment. The EEG data was collected using two EEG systems: Biosemi 64-channels (Biosemi Inc., Amsterdam, The Netherlands) and Neuronaute 21-channels (BioSerenity, Paris, France); 9 participants’ data was recorded using the two systems, respectively, for a total of 18 subjects. Electrode placement for both systems was done according to the 10-20 or extended 10-20 international norm. The data was downsampled to 200Hz.

### Experimental setup and task

Each participant sat comfortably in a chair facing Pepper with 55-60cm distance between them. The Vicon cameras were placed around the participants and Pepper/participant placed at the center. The experimental task was a modified 3D version of the classical Serial reaction time (SRT) Task (Nissen & Bullemer, 1987) inspired by Dominey 1998. It consisted of 9 blocks of 80 hand movements. Participants were instructed to mimic the movement of Pepper as accurately as possible. The blocks B1-4,6,7,9 were training blocks with identical spatial and temporal information within the motor sequence. The training sequence consisted of five repetitions of 16 movements with the sequence -A - - B-C-D - - B-A - - C-D - - C-D - - C - - B-A-D-A- -C - - B-D. The letters represented four different spatial locations. The double dashes represent a temporal pause of 0.5s, and single dashed, no pause. In blocks B5 and B8 we changed either the spatial order of the sequence or the temporal structure of pauses. In the temporal perturbation, the order of the sequential elements was identical to the training blocks but the sequence onset time for each element was systematically manipulated. Conversely, for the spatial perturbation, the elements’ ordering was systematically perturbed, while keeping the sequence timing identical to training blocks. The order of the perturbation type was randomized across participants; thus, 9 participants were presented with a spatial perturbation in B5 and temporal one in B8, while the rest 9 participants received the converse. Throughout the experiment, each block was preceded by 30 seconds of rest, then 30 seconds of fixation on the robot chest screen followed by the sequencing task. After the sequencing, a subsequent fixation period ensued, and then the following rest period for the next block. So, around each experiment block, we had a Rest1 and Fix1 preceding the block and Fix2 and Rest2 following it. In Rest periods, participants were asked to relax and rest their eyes while restricting any unnecessary movements. During the fixation periods, participants had to focus on Pepper’s right-hand finger without making any movements. The fixation period was followed with the experimental sequence learning period where the SRT task is performed followed by Fix2 and Rest2.

### Electrophysiological pre-processing and analyses

#### EEG preprocessing

Electrophysiological (EEG) data for the 18 participants were preprocessed using a customized python based Jupyterlab script. The data was down-sampled to 200 Hz and a band-pass filter (1-40Hz) was applied.. Channel locations were assigned according to the standardized MNI coordinates (BEM dipfit model) over the 21 and 64 channels. Bad channels were removed from data through visual inspection. Independent component analysis (ICA; UNICA format) was applied to the data with a visual artefactual rejection of components with artefacts such as eye blinks, line noise, etc. Event related perturbation spectrum (ERSP) values for theta (3-8Hz), Mu (8-12Hz) and Beta (13-25Hz) were computed. The time-frequency representation of power was computed using Morlet wavelets with three cycles between 0 to 30Hz, and ERSP power values were calculated with a baseline of −600 to −200ms which was expressed in decibels (dB) in 100 log-spaced frequencies. The stimulation-based ERSP power values and time-frequency maps were generated using this customized script for the channels of interest, i.e., Cz, C3, C4, Fz, F3, F4, Pz, P3 and P4. The block onset and offset were synchronized to Pepper’s movement onsets and offsets as performed for the motion capture data.

#### Analyses

Our EEG analyses focused on three aims. First, we aimed to investigate period-specific influences on the ERSP values throughout the experiment. Existing literature suggest a clear desynchronized ERSP state existent during task related movements and a state of synchronisation during periods of rest between tasks (9). To investigate this, 5 Periods (Rest1, Fix1, Movemen,t, Rest2, Fix2) * 3 (theta, mu, beta) frequencies repeated measures ANOVAs was conducted on the ERSP values. The second aim of this paper was to evaluate the motor sequence learning effects using the condition specific ERSP values. For this, repeated measures ANOVAs were conducted to find general learning effects across the 9 blocks and sequence-specific perturbation effects across training and perturbation blocks, respectively. These ANOVAs were conducted on the combined 9 channels of interests (called Grand ERSP from now on) and neural region-specific, i.e., frontal (F3, F4, FZ), central (C3, C4, CZ) and parietal (P3, P4, PZ).

In the general learning effect, the design used was 9 blocks * 3 frequencies whereas the design for sequence-specific effects was 2 (training, perturbation) * 3 frequencies * 2 (spatial, temporal) perturbation type. The training block used in the investigation here was the training block preceding the perturbation block. Finally, while these analyses were performed at the block level, the third objective was to examine at a finer-grained level the relation between information in the EEG signal and the sequential behavior. For this we use a Markov-Switching Linear Regression (MSLR) model in order to discover the underlying structure of the link between the EEG signal and behavior.

### Markov-Switching Linear Regression

Markov-Switching Linear Regression (MSLR) models, which we ran using *Dynamax* (33), are a powerful tool for modeling time series data that exhibit regime-switching behaviour, where the underlying dynamics of the system change over time.

The MSLR model is defined by a set of linear regressions, each associated with a particular state of a discrete Markov chain. The state of the Markov chain determines which sets of weights and biases predicts the evolution of the observed data at each time step. The transitions between states are governed by the transition probabilities of the Markov chain, which are learned from the data.

Formally, an MSLR model can be described as follows. If S is the total number of latent (discrete) states of a Markov process, at each time step t, a given state z_*t*_ (∈ *{*0, 1, …, S*}*) will follow a Markovian evolution such that:

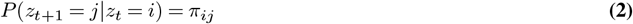

As these are stochastic matrices, π_*ij*_ ∈[0, 1].

Let the *M* −dimensional input time series at time t be denoted by x_*t*_ (∈ ℝ^*M*^). Let the N −dimensional output time series at time t be denoted by y_*t*_ ∈ ℝ^*N*^. Then, in the case of a MSLR, the discrete latent variable at time *t* (z_*t*_), will dictate which emission weights (*W* ∈ ℝ^*N* ×*M*^) and emission biases (b_*s*_ ∈ ℝ^*N*^) we will use to predict the outputs (emissions) based on the inputs (predictors). Moreover, an emission covariance matrix 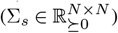 will also have to be learnt. Explicitly, at time t, the emission distribution in this model is given by:

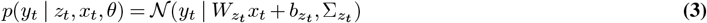

Therefore, the problem of fitting this model amounts to finding the set of emission parameters denoted by:

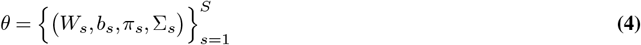

In other words, the aim is to find the weights (W_*s*_) and biases (b_*s*_) for the linear regressions and the transition π_*s*_ and covariance Σ_*s*_ matrices for the Markov process.

In our case, the discrete latent variable (z_*t*_) represents the internal state of the participant at time t, which is inferred from the EEG signal 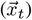 and the observation 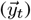 that represents the multi-dimensional output of recorded movement readouts (both from the robot and from the participant). We trained the MSLR model using the Expectation-Maximization (EM) algorithm (34), which iteratively computes the probability over latent states given the data and updates the model parameters to maximize the likelihood of the observed data. For further details, we refer the reader to (35). We iterated the EM algorithm for 50 times, for all models. We initialized the model parameters using a normal distribution for weights and biases and we used the identity matrix as the initial covariance matrix for the emissions. We assumed a Dirichlet prior for the transition matrix. We repeated this process 10 times to increase confidence that we got the optimum value for each combination of parameters.

### Training and inference

We used an 80 : 20 ratio for train-test splitting and performed hyperparameter optimization by cross-validating the training set only (see *Model tuning* for details on CV and model selection). For each species, we concatenated the training sets of all sessions, with forced transitions in between the sessions (setting predictors and emissions to 0 for 50 consecutive trials), so that state probabilities are reset. Then, after optimizing each model, we performed inference on each held out test set (separately per session). We decided to take this approach for various reasons:

1. Model generalization: as the model learns from different participants, it is likely that it can pick up on common information between them.
2. Model interpretability: given that we do not update the model parameters at the inference step, all internal states have the same meaning over participants and, thus, are directly comparable.
3. Better convergence: increasing the number of training samples (i.e. concatenating sessions as opposed to training a different model per session) allows the model to have more data to learn from.

All of the results in the main text, unless otherwise stated, are for held out data.

### Model tuning

For the model we described in *Markov-Switching Linear Regression*, there are several parameters that can be tuned to explain the data better. In our case, we decided to explore the influence of changing the maximum number of internal states (S), to add sticky transitions to the Markov process (a self-bias term in the transition matrix π, making states taking longer to transition to a different one), and to vary the transition matrix sparsity (concentration).

In order to balance model performance with scientific insights, we took a hybrid approach. We increased the number of internal states in a greedy way, to show that the error saturates and that there are diminishing returns when increasing model complexity. On the other hand, for a given number of states, we optimized two free parameters of the Markov process: state stickiness and state concentration. For the sake of efficiency, we used Optuna (36), a flexible framework to implement Bayesian optimization. In Table 1 we report the relevant quantities for this process.

**Table 1.** Parameter values for the Bayesian parameter optimization procedure. These are independently explored for each number of internal states of the HMM.

| Parameter & Value |
| --- |
| Concentration & [0, 100] |
| Stickiness & [0, 100] |
| Sampler & CMA-ES (38) |
| Objective function & $R^2$ |
| Number of Searches & 100 |

To select the best combination of parameters, we performed 5-fold Time-Blocked Cross-Validation (37).

**Fig. S1.**
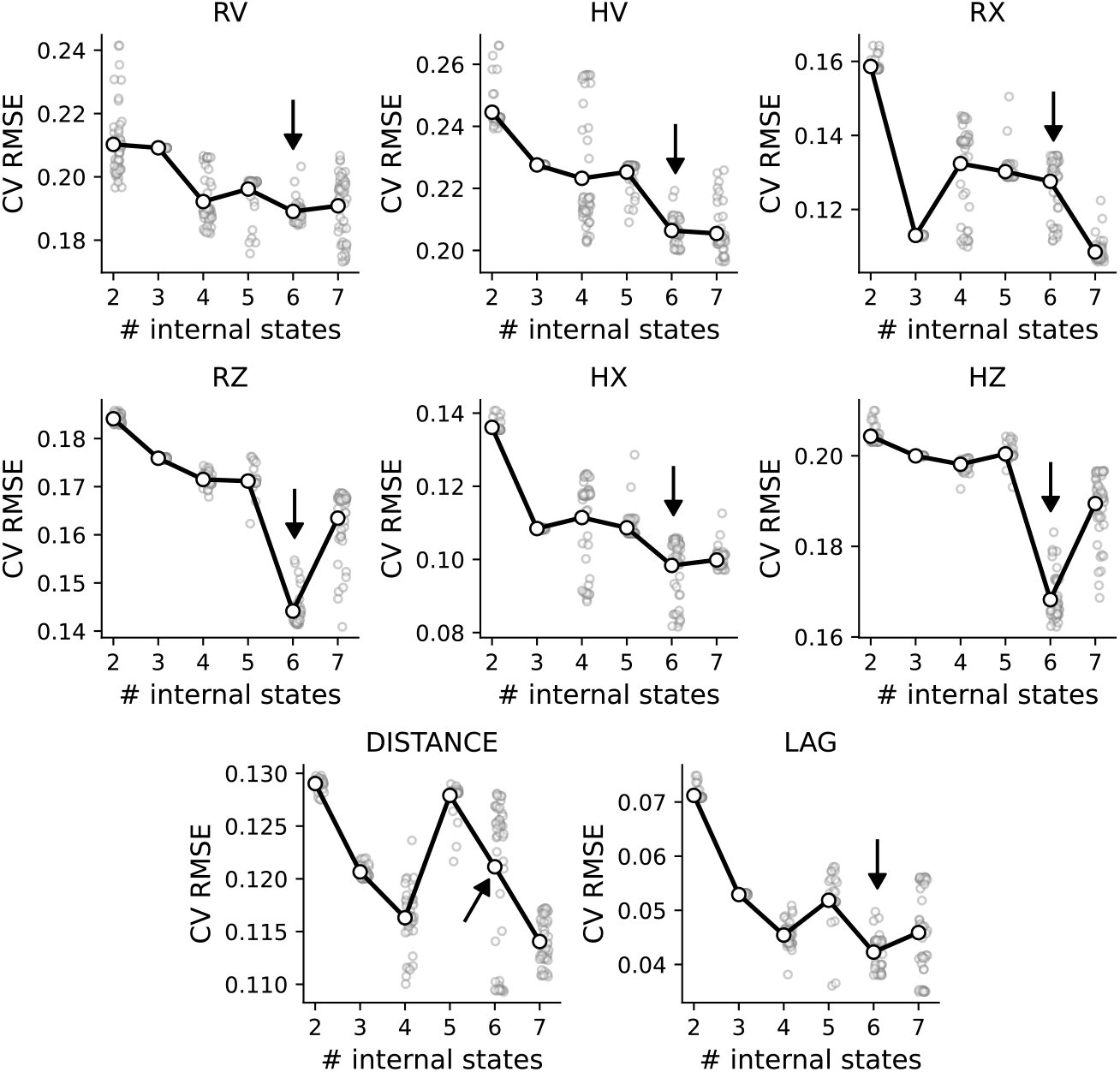
Model selection and optimization. We show the cross-validation performance for a different number of states, for all behavioural readouts. In all of them, there is a clear benefit when allowing the model to switch between more states. The arrows indicate the number of states we ended up selecting (*n*_*s*_ = 6). All readouts are accurately predicted (RMSE ∈ [0.11, 0.18]).

**Fig. S2.**
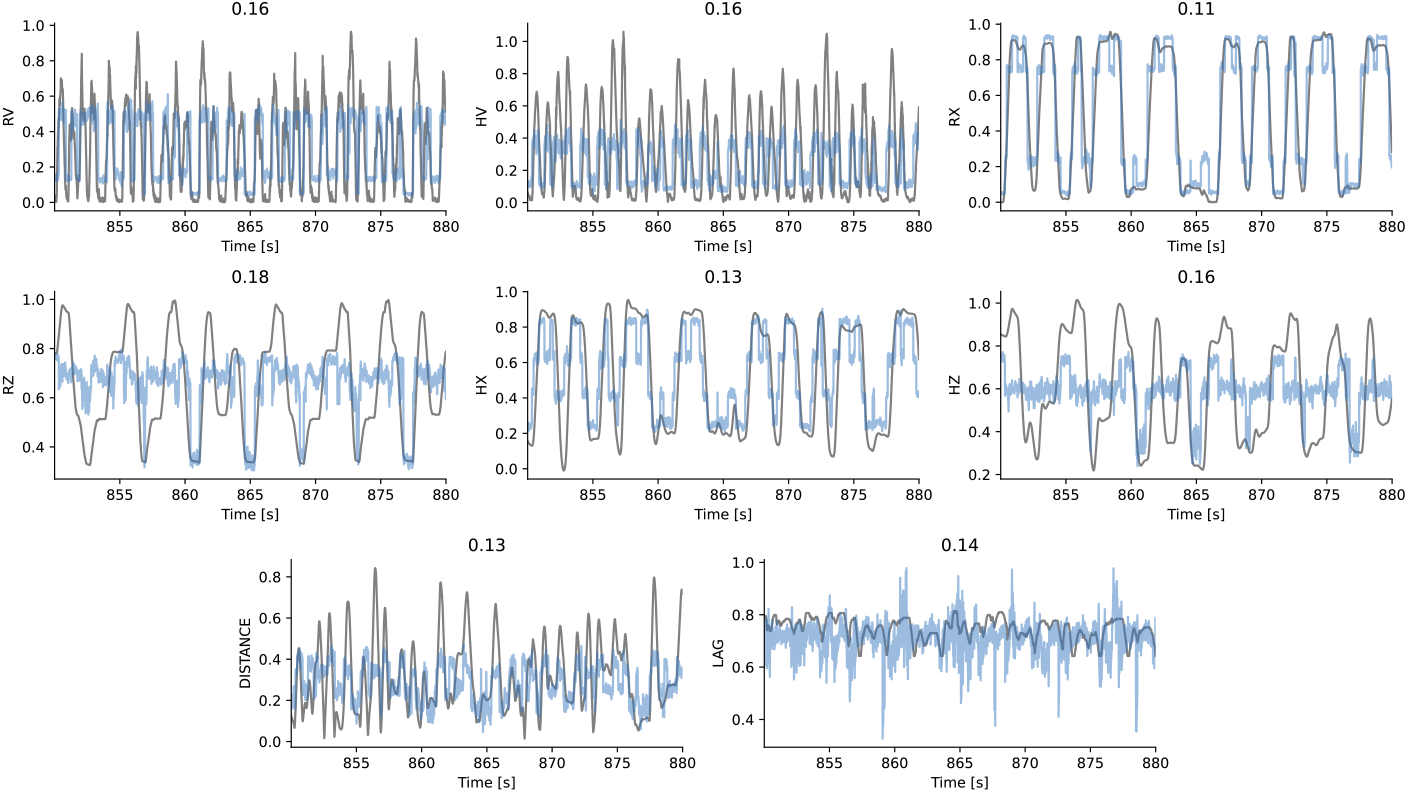
Example measured and predicted traces. We show the ground-truth (gray) and predicted (blue) traces for part of the test set (25 seconds worth of data). As in the Cross Validation case, all readouts are accurately predicted (*RMSE* ∈ [0.11, 0.18]). R-robot, H-human, x,y,z spatial coordinates. Lag- temporal lag between human and robot. Distance - spatial distance between human and robot.

**Fig. S3.**
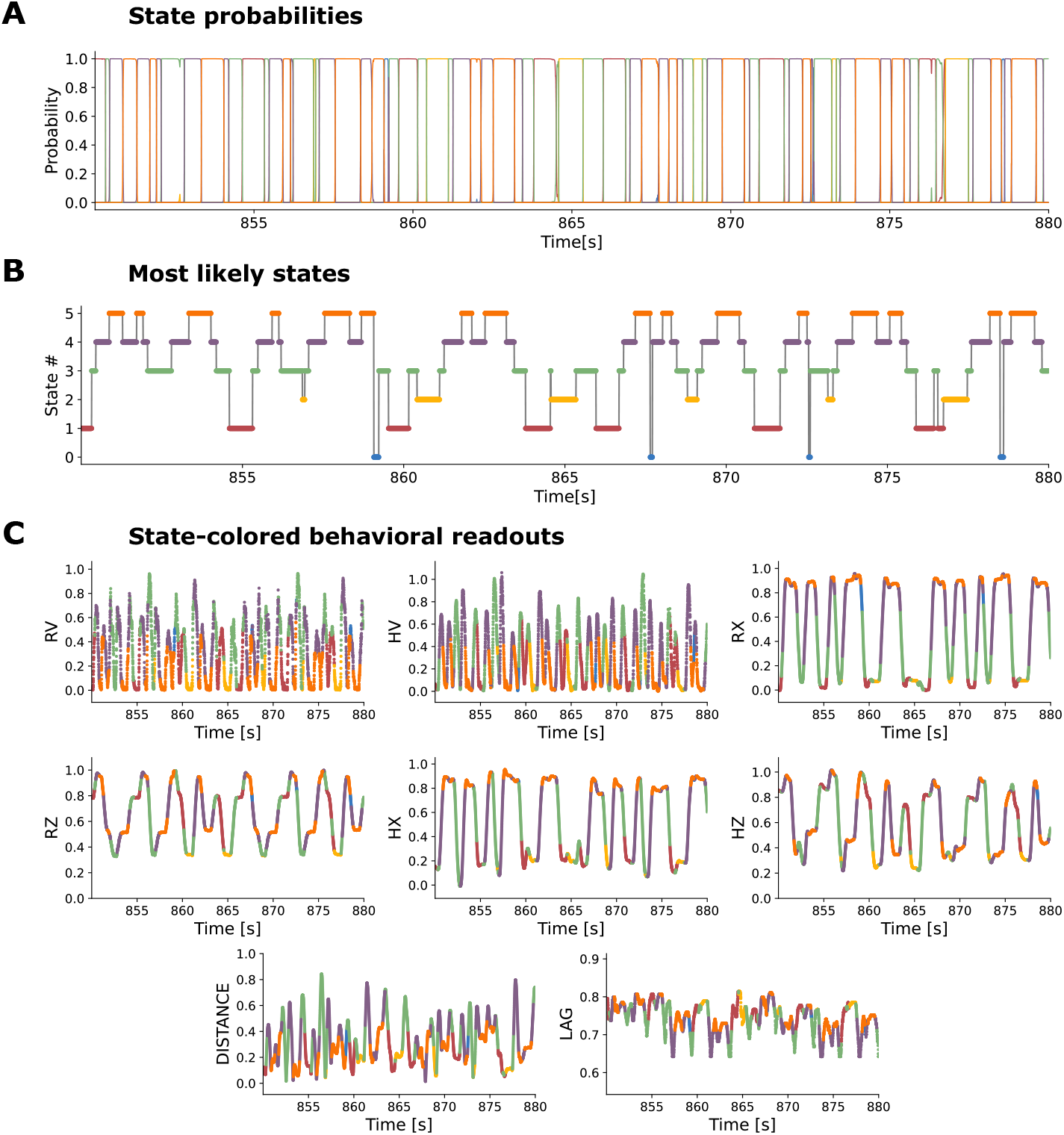
Example state traces.

**Fig. S4.**
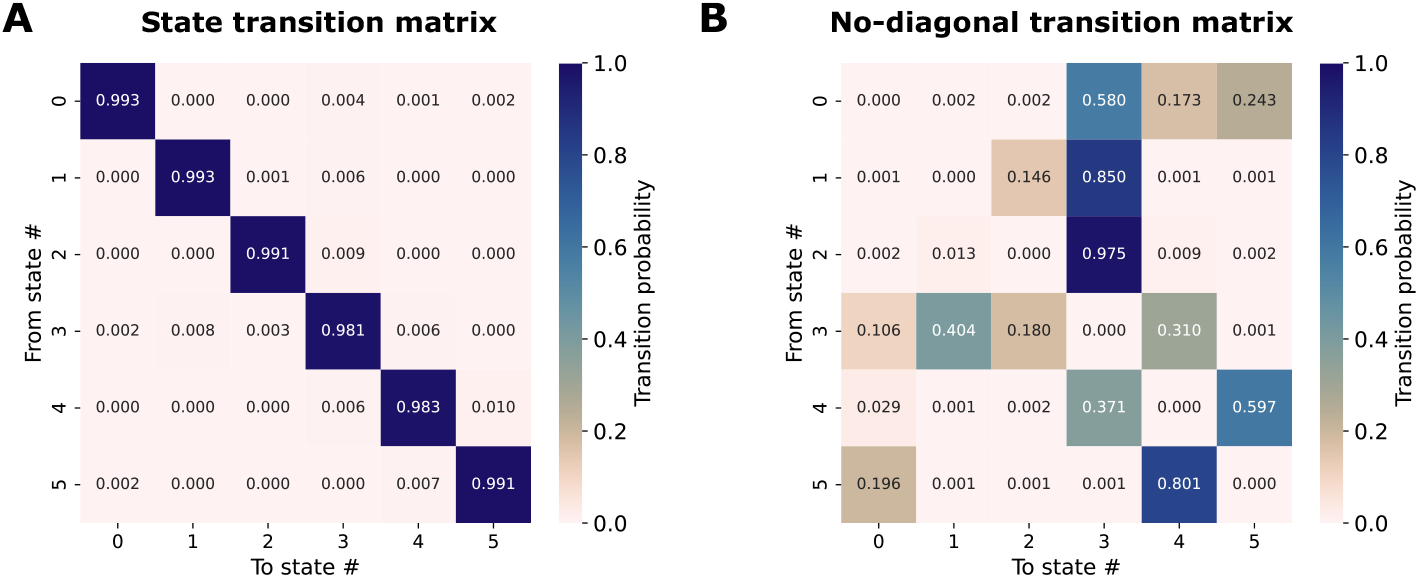
Transition matrix.

**Fig. S5.**
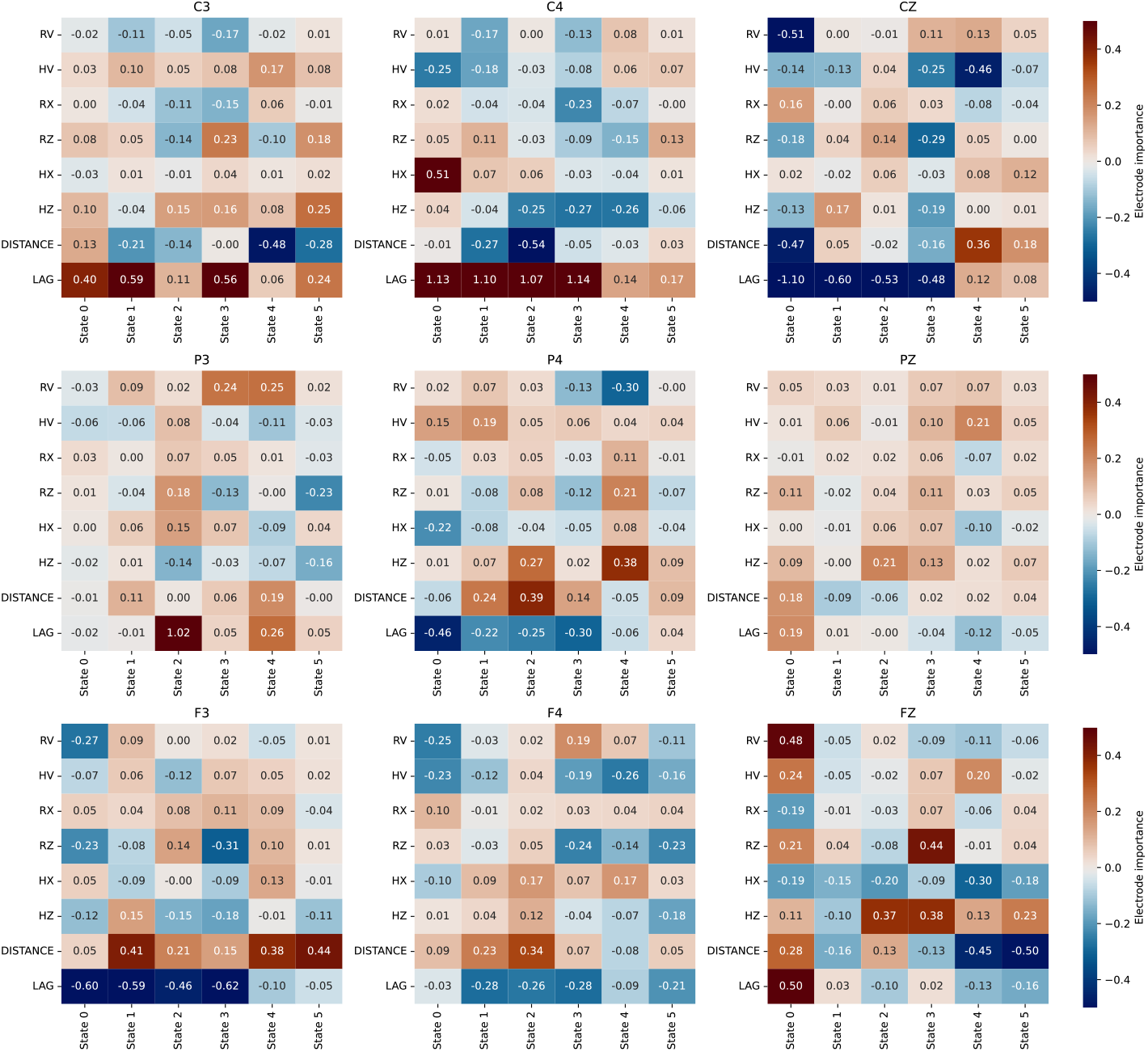
Electrode importance for all readouts, over states. There is a rich structure of variability both over states and over readouts. This implies that the importance of an electrode is dynamical and depends on the movement type.

## Bibliography

1. Karl Spencer Lashley. The problem of serial order in behavior, volume 21. Bobbs-Merrill Oxford, United Kingdom, 1951.

2. Julien Doyon, Ella Gabitov, Shahabeddin Vahdat, Ovidiu Lungu, and Arnaud Boutin. Current issues related to motor sequence learning in humans. Current opinion in behavioral sciences, 20:89–97, 2018.

3. Mary Jo Nissen and Peter Bullemer. Attentional requirements of learning: Evidence from performance measures. Cognitive psychology, 19(1):1–32, 1987. ISSN 0010-0285.

4. Edwin M Robertson. The serial reaction time task: implicit motor skill learning? Journal of Neuroscience, 27(38):10073–10075, 2007.

5. George Kachergis, Floris Berends, Roy De Kleijn, and Bernhard Hommel. Trajectory effects in a novel serial reaction time task. In Proceedings of the Annual Meeting of the Cognitive Science Society, volume 36.

6. Clara Moisello, Domenica Crupi, Eugene Tunik, Angelo Quartarone, Marco Bove, Giulio Tononi, and M Felice Ghilardi. The serial reaction time task revisited: a study on motor sequence learning with an arm-reaching task. Experimental brain research, 194:143–155, 2009. ISSN 0014-4819.

7. P. F. Dominey. A shared system for learning serial and temporal structure of sensori-motor sequences? evidence from simulation and human experiments. Brain Res Cogn Brain Res, 6(3):163–72, 1998. ISSN 0926-6410 (Print) 0926-6410 (Linking).

8. Gert Pfurtscheller. Eeg event-related desynchronization (erd) and synchronization (ers). Electroencephalography and Clinical Neurophysiology, 1(103):26, 1997.

9. Mario Hervault, Pier-Giorgio Zanone, Jean-Christophe Buisson, and Raoul Huys. Cortical sensorimotor activity in the execution and suppression of discrete and rhythmic movements. Scientific Reports, 11(1):22364, 2021. ISSN 2045-2322.

10. Nurhan Erbil and Pekcan Ungan. Changes in the alpha and beta amplitudes of the central eeg during the onset, continuation, and offset of long-duration repetitive hand movements. Brain research, 1169:44–56, 2007.

11. P Zhuang, C Toro, J Grafman, Paolo Manganotti, L Leocani, and M Hallett. Event-related desynchronization (erd) in the alpha frequency during development of implicit and explicit learning. Electroencephalography and clinical neurophysiology, 102(4):374–381, 1997. ISSN 0013-4694.

12. Jarrad AG Lum, Gillian M Clark, Pamela Barhoun, Aron T Hill, Christian Hyde, and Peter H Wilson. Neural basis of implicit motor sequence learning: modulation of cortical power. Psychophysiology, 60(2):e14179, 2023. ISSN 0048-5772.

13. Marzia De Lucia, Irina Constantinescu, Virginie Sterpenich, Gilles Pourtois, Margitta Seeck, and Sophie Schwartz. Decoding sequence learning from single-trial intracranial eeg in humans. PLoS One, 6(12):e28630, 2011. ISSN 1932-6203.

14. Alejandro Tlaie, Muad Y Abd El Hay, Berkutay Mert, Robert Taylor, Pierre-Antoine Ferracci, Katharine Shapcott, Mina Glukhova, Jonathan W Pillow, Martha N Havenith, and Marieke Schölvinck. Thoughtful faces: inferring internal states across species using facial features. bioRxiv, pages 2024–01, 2024.

15. Mikhail Lebedev. Brain-machine interfaces: an overview. Translational Neuroscience, 5:99–110, 2014.

16. Bradley J Edelman, Jianjun Meng, Daniel Suma, Claire Zurn, Eric Nagarajan, Bryan S Baxter, Christopher C Cline, and BJSR He. Noninvasive neuroimaging enhances continuous neural tracking for robotic device control. Science robotics, 4(31):eaaw6844, 2019.

17. Elena Cioffi, Anna Hutber, Rob Molloy, Sarah Murden, Aaron Yurkewich, Adam Kirton, Jean-Pierre Lin, Hortensia Gimeno, and Verity M McClelland. Eeg-based sensorimotor neurofeedback for motor neurorehabilitation in children and adults: a scoping review. Clinical Neurophysiology, 2024.

18. Congzhi Tang, Ting Zhou, Yun Zhang, Runping Yuan, Xianghu Zhao, Ruian Yin, Pengfei Song, Bo Liu, Ruyan Song, Wenli Chen, et al. Bilateral upper limb robot-assisted rehabilitation improves upper limb motor function in stroke patients: a study based on quantitative eeg. European Journal of Medical Research, 28(1):603, 2023.

19. Mohammad Reza Keshtkaran, Andrew R Sedler, Raeed H Chowdhury, Raghav Tandon, Diya Basrai, Sarah L Nguyen, Hansem Sohn, Mehrdad Jazayeri, Lee E Miller, and Chethan Pandarinath. A large-scale neural network training framework for generalized estimation of single-trial population dynamics. Nature Methods, 19(12):1572–1577, 2022.

20. Zoe C Ashwood, Nicholas A Roy, Iris R Stone, International Brain Laboratory, Anne E Urai, Anne K Churchland, Alexandre Pouget, and Jonathan W Pillow. Mice alternate between discrete strategies during perceptual decision-making. Nature Neuroscience, 25(2):201– 212, 2022.

21. Luca Mazzucato. Neural mechanisms underlying the temporal organization of naturalistic animal behavior. Elife, 11:e76577, 2022.

22. Adam J Calhoun, Jonathan W Pillow, and Mala Murthy. Unsupervised identification of the internal states that shape natural behavior. Nature neuroscience, 22(12):2040–2049, 2019.

23. Daniel Hulsey, Kevin Zumwalt, Luca Mazzucato, David A McCormick, and Santiago Jaramillo. Decision-making dynamics are predicted by arousal and uninstructed movements. bioRxiv, pages 2023–03, 2023.

24. Pedro J Gonçalves, Jan-Matthis Lueckmann, Michael Deistler, Marcel Nonnenmacher, Kaan Öcal, Giacomo Bassetto, Chaitanya Chintaluri, William F Podlaski, Sara A Haddad, Tim P Vogels, et al. Training deep neural density estimators to identify mechanistic models of neural dynamics. Elife, 9:e56261, 2020.

25. Byron M Yu, John P Cunningham, Gopal Santhanam, Stephen Ryu, Krishna V Shenoy, and Maneesh Sahani. Gaussian-process factor analysis for low-dimensional single-trial analysis of neural population activity. Advances in neural information processing systems, 21, 2008.

26. Sean M Perkins, John P Cunningham, Qi Wang, and Mark M Churchland. Simple decoding of behavior from a complicated neural manifold. bioRxiv, pages 2023–04, 2023.

27. Matthias Seeger. Gaussian processes for machine learning. International journal of neural systems, 14(02):69–106, 2004.

28. Christopher M Bishop. Pattern recognition and machine learning. Springer google schola, 2:1122–1128, 2006.

29. Luke McEllin, Günther Knoblich, and Natalie Sebanz. Imitation from a joint action perspective. Mind & Language, 33(4):342–354, 2018.

30. Peter E Keller, Giacomo Novembre, and Michael J Hove. Rhythm in joint action: psychological and neurophysiological mechanisms for real-time interpersonal coordination. Philosophical Transactions of the Royal Society B: Biological Sciences, 369(1658):20130394, 2014.

31. Dimitrios Kourtis, Natalie Sebanz, and Guenther Knoblich. Predictive representation of other people’s actions in joint action planning: An eeg study. Social neuroscience, 8(1):31–42, 2013.

32. Natalie Sebanz, Harold Bekkering, and Günther Knoblich. Joint action: bodies and minds moving together. Trends in cognitive sciences, 10(2):70–76, 2006.

33. Chang. P, Harper-Donnelly. G, Kara A, Li X, et al, and Linderman S Murphy K. Dynamax, a library for State Space Models. GitHub, 2022.

34. Arthur P Dempster, Nan M Laird, and Donald B Rubin. Maximum likelihood from incomplete data via the em algorithm. Journal of the royal statistical society: series B (methodological), 39(1):1–22, 1977.

35. Scott Linderman, Matthew Johnson, Andrew Miller, Ryan Adams, David Blei, and Liam Paninski. Bayesian learning and inference in recurrent switching linear dynamical systems. In Artificial Intelligence and Statistics, pages 914–922. PMLR, 2017.

36. Takuya Akiba, Shotaro Sano, Toshihiko Yanase, Takeru Ohta, and Masanori Koyama. Optuna: A next-generation hyperparameter optimization framework. In Proceedings of the 25th ACM SIGKDD international conference on knowledge discovery & data mining, pages 2623–2631, 2019.

37. Tom AB Snijders. On cross-validation for predictor evaluation in time series. In On Model Uncertainty and its Statistical Implications: Proceedings of a Workshop, Held in Groningen, The Netherlands, September 25–26, 1986, pages 56–69. Springer, 1988.

38. Nikolaus Hansen. The cma evolution strategy: A tutorial. arXiv preprint 1604.00772, 2016.

